# Malaria in Southern Venezuela: The Hottest Hotspot in Latin America

**DOI:** 10.1101/2020.03.13.990457

**Authors:** ME Grillet, JE Moreno, JV Hernández, MF Vincenti-González, O Noya, A Tami, A Paniz-Mondolfi, M Llewellyn, R Lowe, AA Escalante, JE Conn

## Abstract

Malaria cases in Latin America reached ~1 million in 2017 and 2018, with 53% and 51% reported from Venezuela, respectively. In this study, we characterized the spatiotemporal dynamics of malaria transmission between 2007-2017 in southern Venezuela, the main endemic area of the country. We found that disease transmission was focal and more prevalent in the southeast of southern Venezuela where two persistent hotspots of *Plasmodium vivax* (76%) and *P. falciparum* (18%) linked to deforestation for illegal gold mining accounted for ~60% of the country-wide number of cases. Incidence has increased nearly tenfold in the last decade, showing an explosive epidemic growth due to a significant lack of disease control. We suggest that a source-sink pattern of *Plasmodium sp*. dispersal account for the re-emergence and progression of malaria transmission in the last 4 years across the country due to the internal migration of infected people to and from the hotspots and other malaria-prone ecosystems. We observe a similar pattern explaining the spillover of cases across international borders affecting neighboring countries. This study provides baseline epidemiological data and guidance for malaria control to further assess the dynamics of cross-border malaria, the role of asymptomatic carriers, drug-resistant evolution, and innovative control efforts in the Latin America region.

## Background

Malaria control programs in the Americas are facing obstacles that are threatening the global strategy for malaria elimination set for the region for 2030 [1,2,3]. Progress in reducing malaria morbidity and mortality between 2000-2015 stalled in 2016 and reversed in 2017 and 2018 [3,4,5] when cases climbed to ~1 million in both years [2,3]. This surge was mainly driven by the collapse of the Venezuelan economy since 2014, with the consequent crumble of its entire healthcare system [6,7]. In particular, Venezuela reported a total of 1,255,299 cases between 2015-2018, the highest number in Latin America, and in 2017 exhibited one of the most substantial increases in malaria cases worldwide [1,2,8]. As in other endemic countries in South America [5], *Plasmodium vivax* accounts for 76 % of reported cases in Venezuela followed by *Plasmodium falciparum* (17.7 %), mixed *P. vivax*/*P. falciparum* infections (6%) and *Plasmodium malariae* <1% [3]. After the successful elimination of malaria in approximately 75% of its territory during the early 1960s [9], low to moderate malaria transmission by *P. vivax* and *P. falciparum* persisted in Venezuela in the lowland Amazon rainforests and savannas of the remote Guayana region, mainly among isolated Amerindians communities, south of the Orinoco River [10]. During the 1980s, *P. vivax* malaria reemerged along the coastal wetlands in the northeastern region of the country [11], but transmission was interrupted twenty years later [12]. However, Bolívar state, a region in the southeast of Venezuela bordering with Brazil and Guyana has been a persistent focus of malaria transmission. Specifically, from 1990 onwards, the principal focus of malaria in Venezuela has been located in this region, contributing > 60% (1992-1995) to 88% (2000-2014) of the country’s total malaria cases [12,13]. The health situation in this region has worsened significantly in recent years, and the current limitations in malaria surveillance and control are a cause of major concern [7,13]. Additionally, since 2014, local malaria transmission has re-emerged in previously endemic and new areas across the whole country, including the northeastern region of Venezuela [7].

At present, the uncontrolled upsurge of malaria compounded by the mass migration of Venezuelans to neighboring countries pose a serious threat to the wider region [6,7,14], jeopardizing the efforts of bordering malaria-endemic countries to achieve their goals for disease control. Human population movement among the countries that form the Guiana Shield (Guyana, French Guiana, Suriname, as well as parts of Colombia, Venezuela, and Brazil) serves to amplify the regional impact of the malaria increase in Bolívar state. There is a genuine risk that a regional malaria corridor will form [15] from Bolívar state to northern Brazil *via* the mass migration of displaced individuals who take advantage of the existing road network (Route 10). This situation is worsened by the presence of novel mutations linked to artemisinin resistant due to irregular malaria treatment in Guyana [16], raising concerns about the potential regional spread of mutations linked to drug-resistant and artemisinin delayed parasite clearance in *P. falciparum* where Venezuela could act as a hub.

Considering all these factors, a first step in understanding how to manage this public health crisis is to characterize the spatiotemporal dynamics of malaria transmission in southern Venezuela. Thus, we analyzed reported malaria cases data (*P. vivax* and *P. falciparum*) between 2007 and 2017 in Bolivar state to identify hotspots, to determine their transient or persistence dynamics, the factors driving them, and their role in the malaria surge observed in the last years across the whole country. Particularly, we tested the hypothesis that illegal gold mining activities disproportionally contribute to the transmission of both parasites in the detected hotspots, a socioeconomic risk factor that has been suggested in previous studies [7,13,17]. Finally, we described the spatial epidemiology of malaria at the national level and assess the extent to which landscape epidemiology of this infection has changed during the 2014-2017 period.

## Study area and Methodology

### Materials and Methods

#### Study area

Venezuela is located on the northern coast of South America and has a surface area of ~916,445 Km^2^ (Fig. 1a). The country has vast regions suitable for malaria transmission with a humid tropical climate modified by altitude and semi-annual seasonal rainfall cycles. Most of the annual precipitation (88%) occurs from April-November (rainy season), followed by a drier season from December-March. Mean daily temperatures are around 28°C throughout most of the country. Bolívar state (Fig. 1b), the focus of this study, covers 240,500 km² and as of the 2018 Venezuelan census [18], has a population of 1,837,485 inhabitants heterogeneously distributed in 11 municipalities and 46 parishes (municipalities in Venezuela are divided into parishes, a third-level administrative unit with specific functions, including reporting health statistics of transmissible diseases). Bolivar state has a low average population density (5.1 inhabitants per km^2^) and extensive territory. Population densities per municipality range from 553 (e.g., Caroni municipality) to < 1 inhabitant per km^2^ (e.g., Gran Sabana municipality). Most of the population resides in the northernmost and easternmost parts of the state (Fig. 1b), where economic activities are related to mining (iron, gold, bauxite, and diamonds), production of steel, aluminum, and hydroelectric industries, businesses and services, forestry, cattle, and agricultural development. The state capital is Ciudad Bolivar (Heres municipality) but Ciudad Guayana (Caroni municipality) is the main urban center. Although these northern cities are well-connected with the rest of Venezuela, the rural southern and western areas are sparsely populated and have few or no roads. There is one major paved road, Route 10, along the eastern side of Bolívar that connects Venezuela with Brazil, crossing the main endemic malaria region in the state (Fig. 1b).

**Figure 1.**
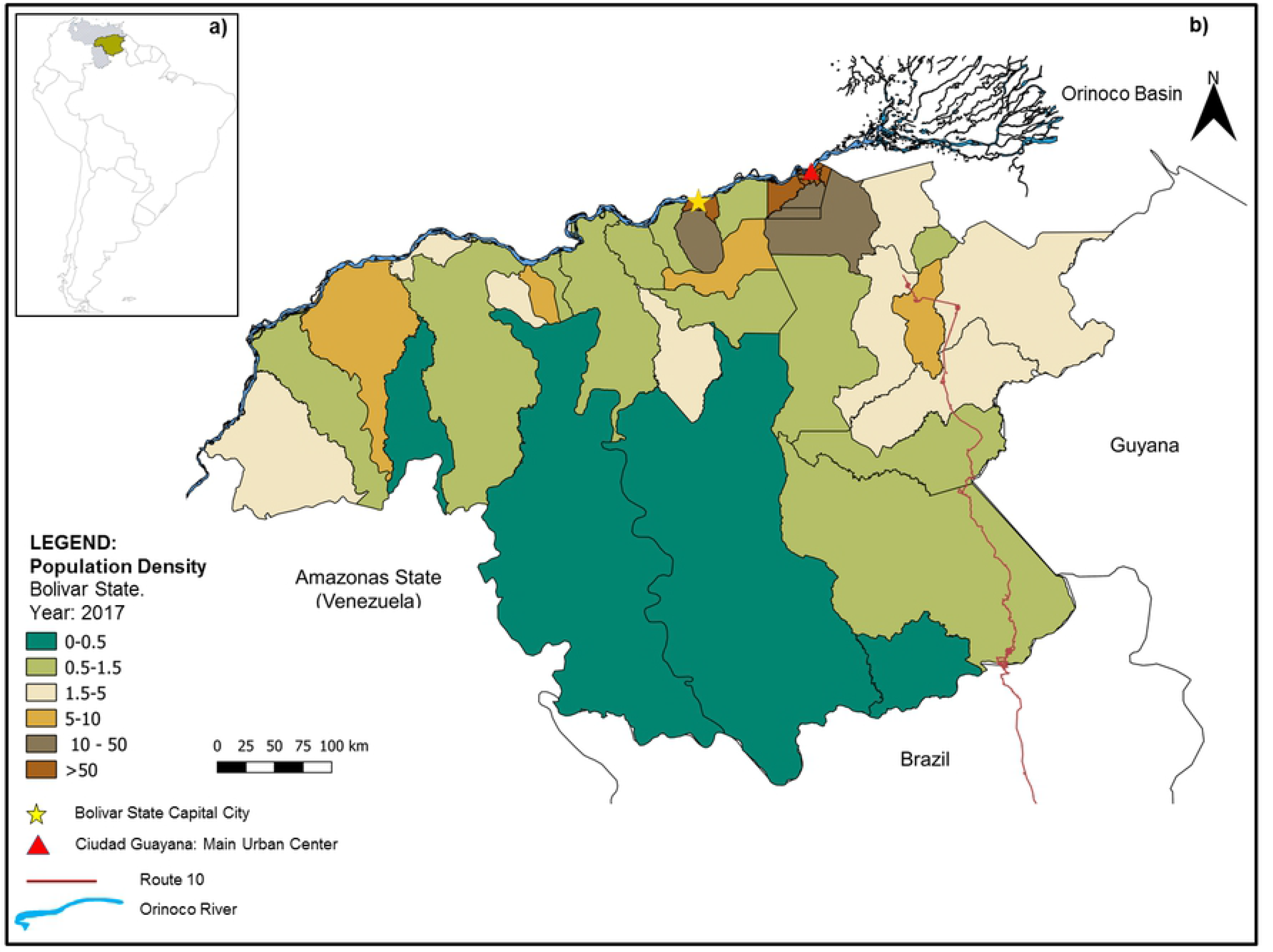
a) Map of Venezuela in South America and the study area (Bolívar state in green). b) Map of Bolívar state showing the population density by civil parish (people per square kilometre), and others relevant features of the landscape.

Regarding Bolívar state, here the lowland malaria areas are characterized by an average temperature of 24-26°C and 1,000-1,500 mm annual rainfall. Rains peak from May to July, followed by a secondary but lower increase in rainfall from October to November. However, malaria in this region has a less marked seasonality, especially *P. vivax*, characterized by a near perennial incidence [12,13]. The primary vectors for all *Plasmodium* species in the state are *Nyssorhynchus darlingi* (also referred to as *Anopheles darlingi*) and *Ny*. *albitarsis s.l*. (also referred to as *Anopheles albitarsis*) [17,19,20,21].

#### Epidemiological and Sociodemographic Data

The yearly number of *P. vivax* and *P. falciparum* cases per parish (2007-2017) in Bolivar state were provided by the Malaria Control Program, a local branch of the Venezuelan Ministry of Health [22]. Time series of annual malaria incidence (2014-2017) per municipality for Venezuela and the whole country were obtained from the PAHO Malaria Surveillance Indicators [23]. Malaria incidence rates per 1,000 inhabitants by parish and by *Plasmodium* species were calculated. These rates were calculated by taking into account the human population growth rate predicted for the studied period according to the demographic data (at-risk population) from the National Statistics Office of Venezuela [18]. In addition, we obtained information about the occupation of individuals presenting with malaria [22]. This allowed us to investigate whether or not miners were at greater risk of malaria in hotspot areas of Bolívar state. We also examined potential associated or confounding factors, including age, sex, and source of infection.

#### Data Analyses

The annual malaria incidence per parish in Bolivar was mapped to characterize spatio-temporal disease dynamics. Local spatial autocorrelation analysis was performed using Anselin Local Moran’s *I* test to identify significant spatial patterns of malaria [24]. Specifically, we evaluated the likelihood of malaria occurring equally at any location (parish centroid) within Bolivar state vs. the detection of unusual aggregations of malaria incidence. This test evaluates adjacent positions and finds a strong positive spatial autocorrelation when the surrounding incidence of disease over the entire study area has analogous values (called “High-High” areas). A positive value for ‘*I*’ indicates that the adjacent parishes are bounded by *P. vivax* or *P. falciparum* incidences with similar values. Such a feature is considered a cluster or hotspot. To test whether the observed clustering/dispersing is statistically significant, an estimated *Z*-score and a 99% level of significance (*P* < 0.001) were selected.

For the detected hotspots, we estimated *R*_0_ or the effective reproductive ratio, *R_E_*, for each parasite species during the last 4 years (2014-2017) due to that was the period when malaria increased exponentially as it will be showed later on. The effective reproductive ratio for malaria is the expected number of secondary cases of *P. vivax* (*PvR*_0_) or *P. falciparum* (*PfR*_0_) that arise from one primary case in a partially immune population (*R_E_* = *sR*_0_) during the infectious period, where *s* is the fraction of the population that is susceptible. Here, we assumed that during the initial spread phase of the incidence epidemic curve in each seasonal (annual) period, susceptible exhaustion of individuals, *s*, is negligible. Consequently, we fitted an exponential growth continuous function to the reported curve of accumulated incidence (log) per week and then estimated *R_0_* as the calculus of the rate of exponential growth *r* of the initial period of the exponential growth function. Therefore, *R_0_* = *Vr*+1 [25], where *V* is the serial interval, *i.e*., the mean time between infection and reinfection in one individual. In one serial interval, each infected individual is assumed to give rise to *R_0_* secondary cases and one removal [25]. The ‘one’ in this expression means that the calculus of *R*_0_ reflects the total number of new infections, whereas the overall growth rate *r* includes the death of the founding individual [26]. Here, we chose mean values of *V* as 30-90 days for *P. vivax* and 30 days for *P. falciparum*.

We explored occupational risks in hotspot areas by analyzing the proportion of the population that worked primarily in mining, agriculture, forestry, fishing, retail, and other, as well as examining the ratio of male-to-female and age-structure from the surveyed population for malaria in Bolivar state. The chi-squared test allowed us to determine whether there were significant differences between the expected frequencies and the observed frequencies in one or more categories. Additionally, to identify illegal mining operations in the study area, we obtained data on the deforested land cover distribution areas in the parishes classified as hotspots across the study period via Global Forest Watch [27] (http://www.globalforestwatch.org) and the visual interpretation of remote sensing images from Landsat Thematic Mapper 8 imagery [28] (https://landsat.gsfc.nasa.gov/landsat-data-continuity-mission/).

Finally, we used kriging, a local geostatistical interpolation method [29], to generate an estimated continuous surface from the scattered set of points (i.e., municipality centroids) to better capture the local spatial variation of malaria spread across the country during the 2014-2017 period. We used an ordinary kriging model to predict the values of the annual parasite incidence (API) at the country level (spatial risk diffusion map) and to estimate the associated errors at unmeasured locations. Several variogram models were tested using a cross-validation procedure. The best model was adjusted for any directional spatial trend in our data (anisotropy) in the semi-variogram [29].

Maps, hotspots, remote sensing image visualization, and kriging analyses were performed in ArcGIS desktop software (version 10.5.1 Redlands, CA: Environmental Systems Research Institute). The calculus of *Ro* was estimated using the ‘epimdr-package’ developed by Bjørnstad [25] and run in R-software (The R-Development Core Team, http://www.r-project.org).

## Results

The total number of accumulated cases of malaria in Venezuela during the sampled decade was 1,207,348 (range: 32,037-411,586), with overall malaria incidence rates (cases/1,000 inhabitants-year) increasing from 5.2 (2007) to 28 (2017; Fig. 2). During that period malaria incidence multiplied nearly 10-fold (2007: 41,749 cases; 2017: 411,586) with a sharp (exponential) and significant increase since 2014 (*R^2^* = 0.92, *P* < 0.05). The mortality due to malaria infection increased nearly 20-fold from 16 (2007) to 312 cases in 2017. Bolívar contributed about half (~47%) of the total cases in Venezuela during 2017, but this proportion represented from 60 to 80% in the previous years, demonstrating that malaria reported from the whole Venezuela in the last decade has essentially tracked that observed in the southeastern region (Fig. 2). In Bolívar, 20-30% of malaria cases were due to *P. falciparum*, the rest (70-80%) were attributed to *P. vivax*, with a mean ratio of *P. vivax*/*P. falciparum* of 3.04 (+/− 0.17 SE). Figure 2 also reveals that the malaria burden due to *P. vivax* in Bolívar State has had a similar and steady 4.5-fold increment (*R^2^* = 0.93, *P* < 0.05) since 2014 (from 32,791 to 146,885 cases) with an accumulated 496,847 cases for the period. A more moderate increase (~2.5-fold, *R^2^* = 0.89, *P* < 0.05) was detected for malaria morbidity due to *P. falciparum* (18,127 cases in 2014 to 43,152 in 2017), with 152,589 accumulated cases for the same period (Fig. 2).

**Figure 2.**
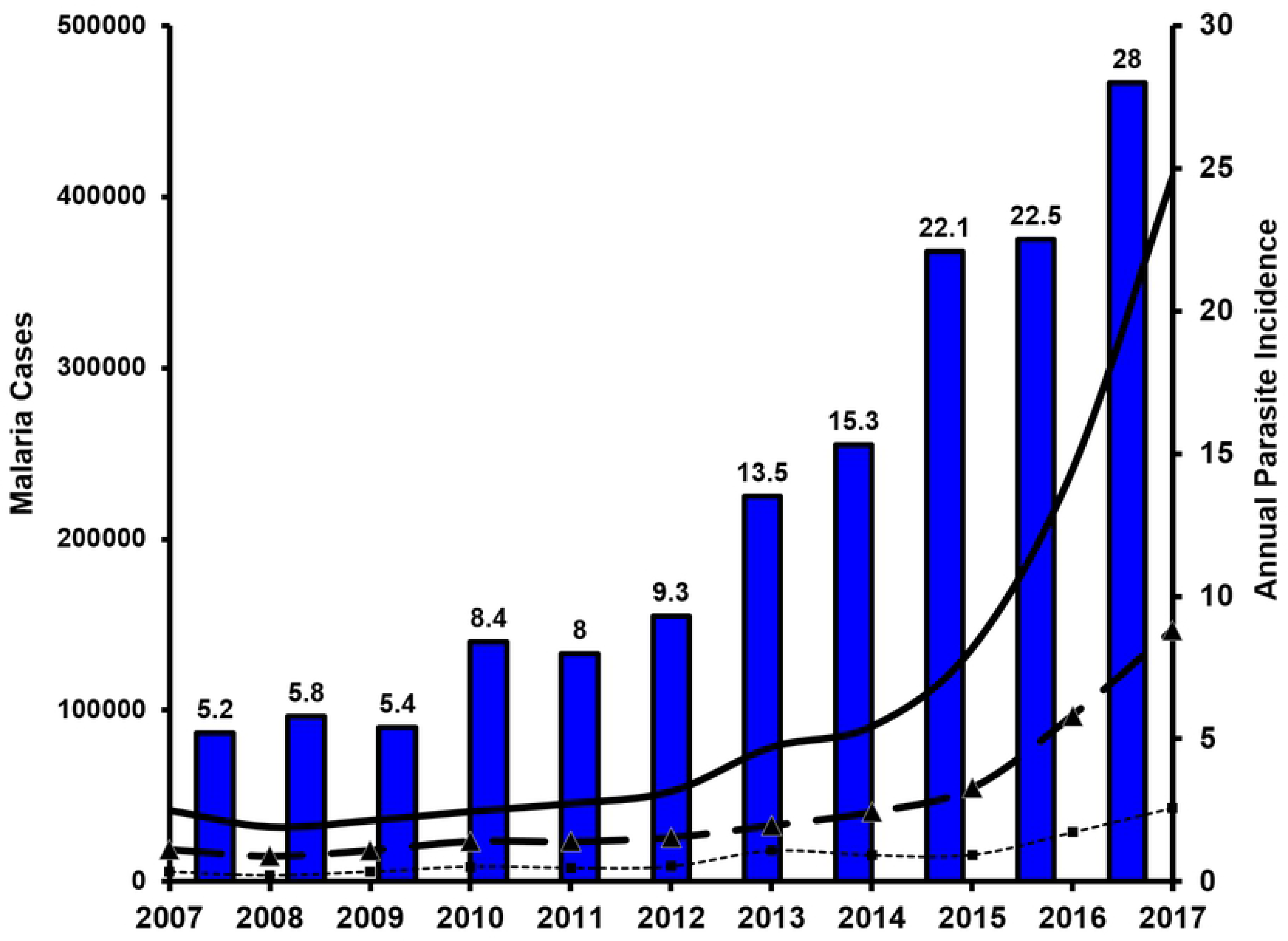
Number of confirmed malaria cases in Venezuela (solid line) and Bolívar state (dashed line: *P. vivax* and small dashed line: *P. falciparum*). Annual parasite incidence (API: Number of confirmed malaria cases/1,000 inhabitants, blue bars) in Venezuela (2007-2017).

At the scale of Bolívar state, annual *P. vivax* incidence from 2007-2017 (Fig. 3) was heterogeneously distributed with most cases spatially concentrated in the mid-eastern and southern parishes of San Isidro, Dalla Costa and Ikabarú (the latter bordering Brazil), followed by northern Pedro Cova, Santa Barbara, and El Callao Parishes. Few or no cases were reported in the remaining parishes. Positive parishes had annual *P. vivax* incidences as high as 4,672 cases/1,000 inhabitants (2017) and mean annual incidences of up to 1,919 cases/1,000 inhabitants-year (e.g., San Isidro). The temporal dynamics of *P. vivax* cases in San Isidro were representative of the dynamics of malaria overall in Bolívar state (data not shown) accounting for ~43.4% of the malaria morbidity during the study period. As with *P. vivax*, spatial heterogeneity was also detected for *P. falciparum* (Fig. 4). In this parasite species, the highest annual incidence was 1,549 cases/1,000 inhabitants (2017) and the highest annual mean was 738 cases /1,000 inhabitants (2017), both also reported for San Isidro Parish.

**Figure 3.**
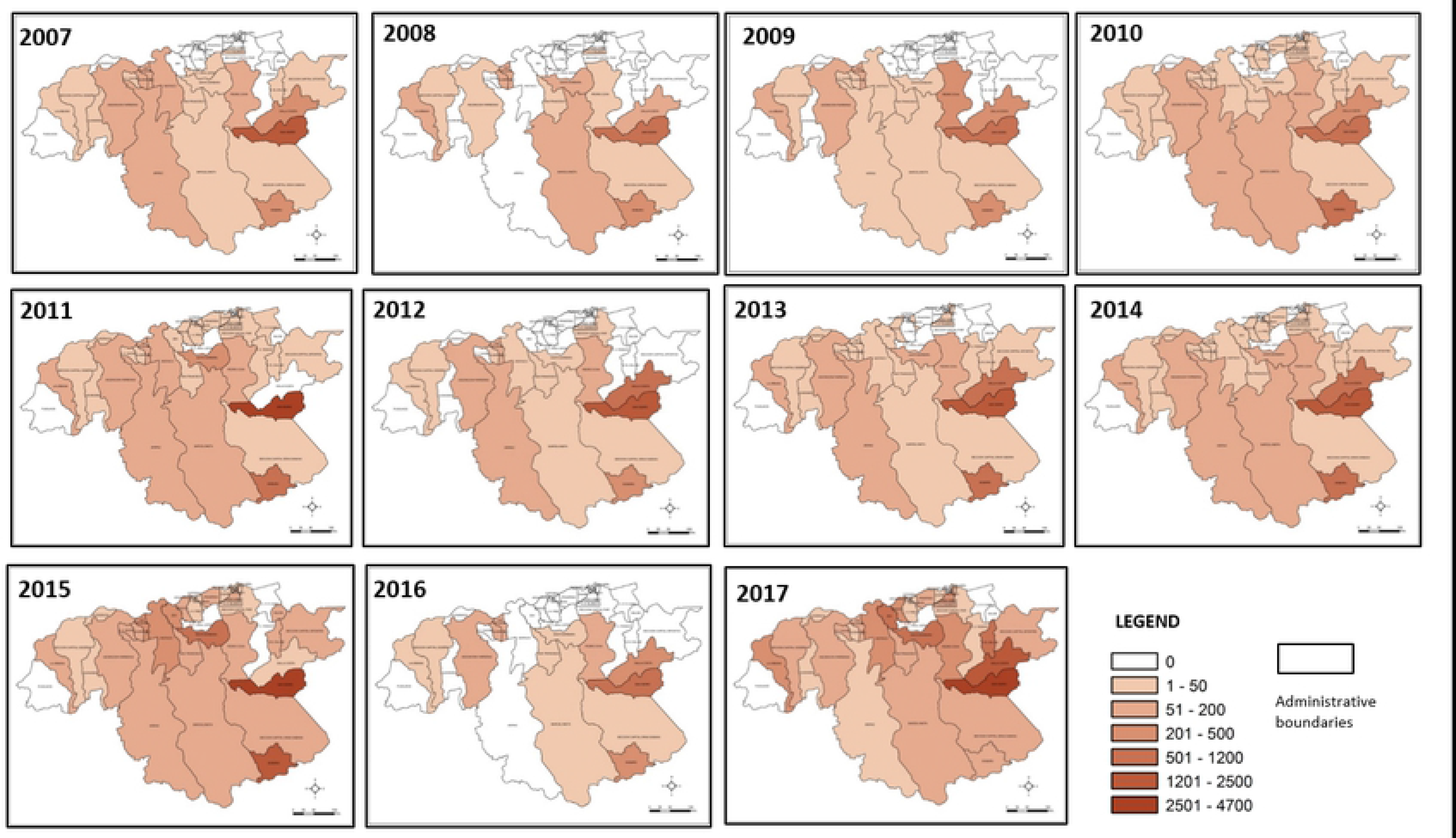
Annual *P. vivax* incidence per parish in Bolívar state (2007-2017), south-eastern Venezuela.

**Figure 4.**
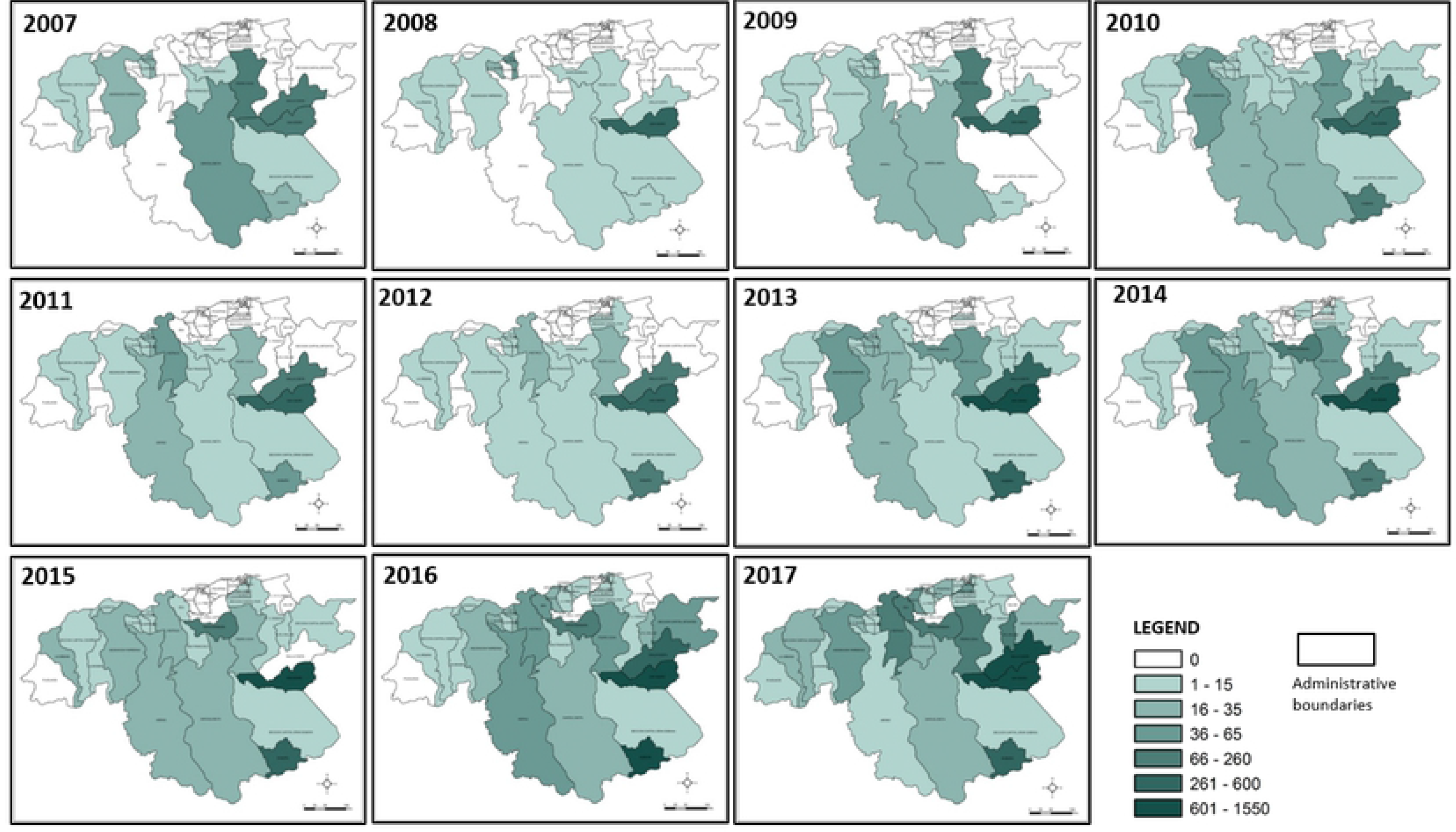
Annual *P. falciparum* incidence per parish in Bolívar state (2007-2017), south-eastern Venezuela.

Additional spatial analyses indicated that significant local clustering of *P. vivax* and *P. falciparum* incidences were detected in the two parishes of San Isidro and Dalla Costa, providing further evidence that malaria transmission has focused and persisted in these localities supporting their ‘hotspot’ status in Bolívar state over the last decade (Figs. 5 and 6). A few exceptions to this pattern were observed for *P. falciparum* during the first three years of the study period (Fig. 6) and for *P. vivax* in 2007 (Fig. 5).

**Figure 5.**
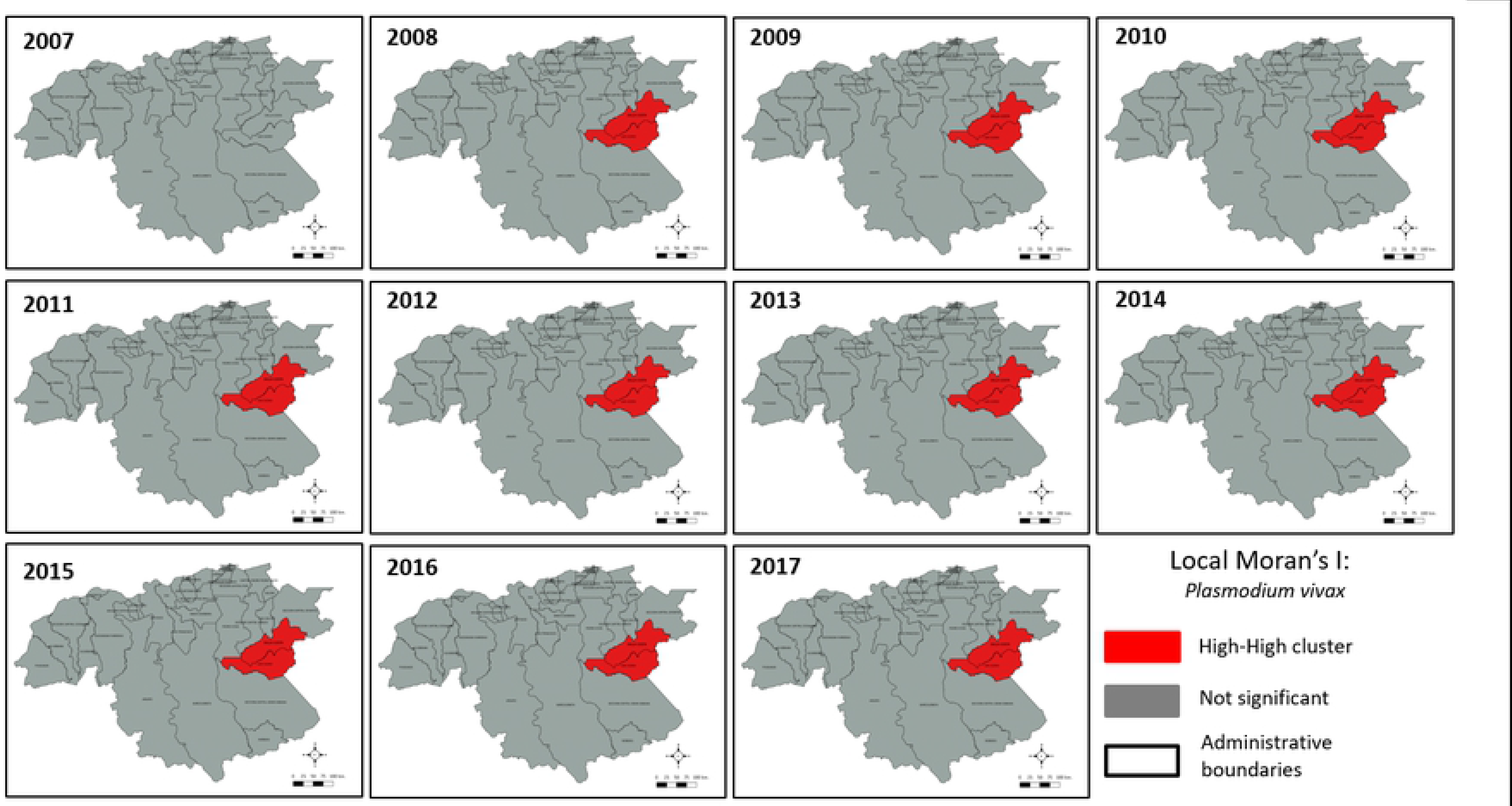
Significant clusters (hotspots) of annual *Plasmodium vivax incidence (*red-red) in Bolívar state (south-eastern Venezuela).

**Figure 6.**
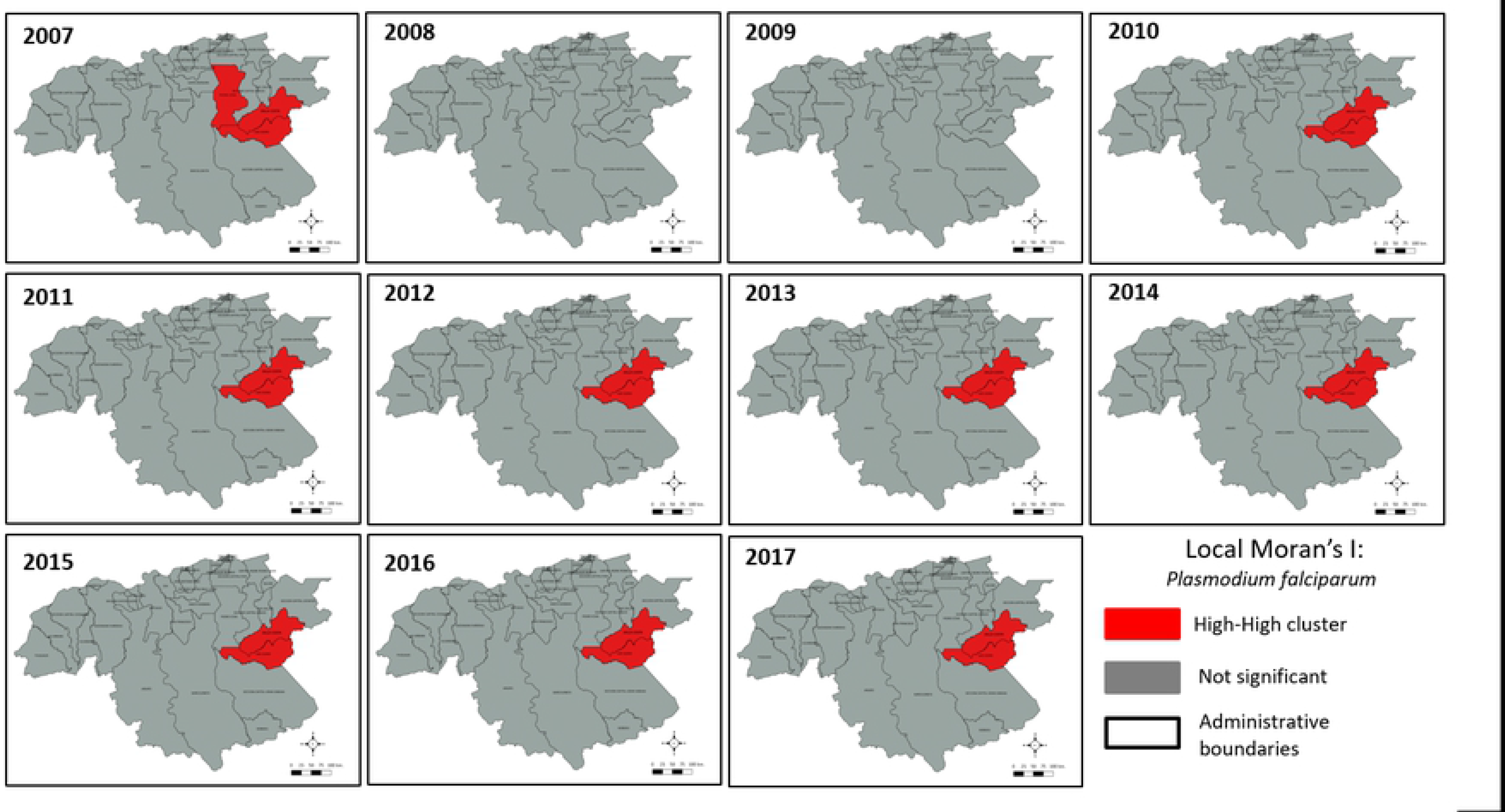
Significant clusters (hotspots) of annual *Plasmodium falciparum incidence (*red-red) in Bolívar state (south-eastern Venezuela).

From the *P. vivax* time-series in San Isidro parish, we estimated values of the reproductive ratio (*PvR*_o_), that ranged from 1.23 - 2.27 over the past four years (S1 Table). Lower estimates (< 2) *R_0_* for *P. falciparum* (*PfR*_o_) were found (S1 Table). Similarly, lower values of *R_o_* were obtained from the epidemic curves for both parasites in Dalla Costa parish (S1 Table).

In San Isidro, across the study period *P. vivax* cases occurred predominantly in men (70%: 145,492/206,455; *X*^2^=34,608.48, 1*df*, *P* < 0.0001), and mainly affected the age-groups of 21-30 (32%), 11-20 (22%) and 31-40 (20%) years (S1 Figure). Occupationally, 62% (127,898/206,455; range: 56%-68%; *X*^2^=11,792.08, 1*df*, *P* < 0.0001) of the surveyed population with *P. vivax* was significantly associated with gold mining activities (miners) compared with other common occupations including housekeeper, machine operator, driver, transporter, student, and agriculture. A similar significant association was found for *P. falciparum*: individuals with gold mining as occupation accounted for 66% of cases (50,013/75,124; range: 63%-70%; *X*^2^=8,254.48, 1*df*, *P* < 0.0001).

The concentration of malaria cases in southern San Isidro coincided with deforested areas resulting directly from illegal mining activities. During the study period, most localities with >1,000 cases in Sifontes municipality (San Isidro and Dalla Costa parishes) were located in this open (deforested) area (Fig. 7a), where the percentage of deforestation (tree cover loss) has dramatically increased in recent years (Fig. 7b). Since 2007, San Isidro concurrently lost 3,058 Ha tree cover (~1.02% decrease) while malaria increased by ~746 % (Fig. 8a). Similarly, Ikabaru Parish (municipality of Gran Sabana), lost 2,934 Ha of tree cover (~1.06% decrease) and registered a significant and parallel malaria increase (Fig. 8b). Figure 9a-d depicts both the spatial spread of malaria cases across Venezuela that has expanded from southern Guayana toward the northern-central-western areas during 2014-2017 (Epidemiological week-EW 21) and the intensification of disease transmission in the south, an endemic area of continued concern. Particularly, this spatial prediction analyses emphasizes that the primary high-risk malaria areas and potential sources of parasite dispersal within the country are the Bolívar state hotspots, followed by the southwestern Amazonas state. Considering the population growth during that period, the national percentage of individuals living in areas at risk of contracting malaria increased from 34.4 % (9,907,708 people) to 50 % (15,988,534 people) between 2014 and 2017.

**Figure 7.**
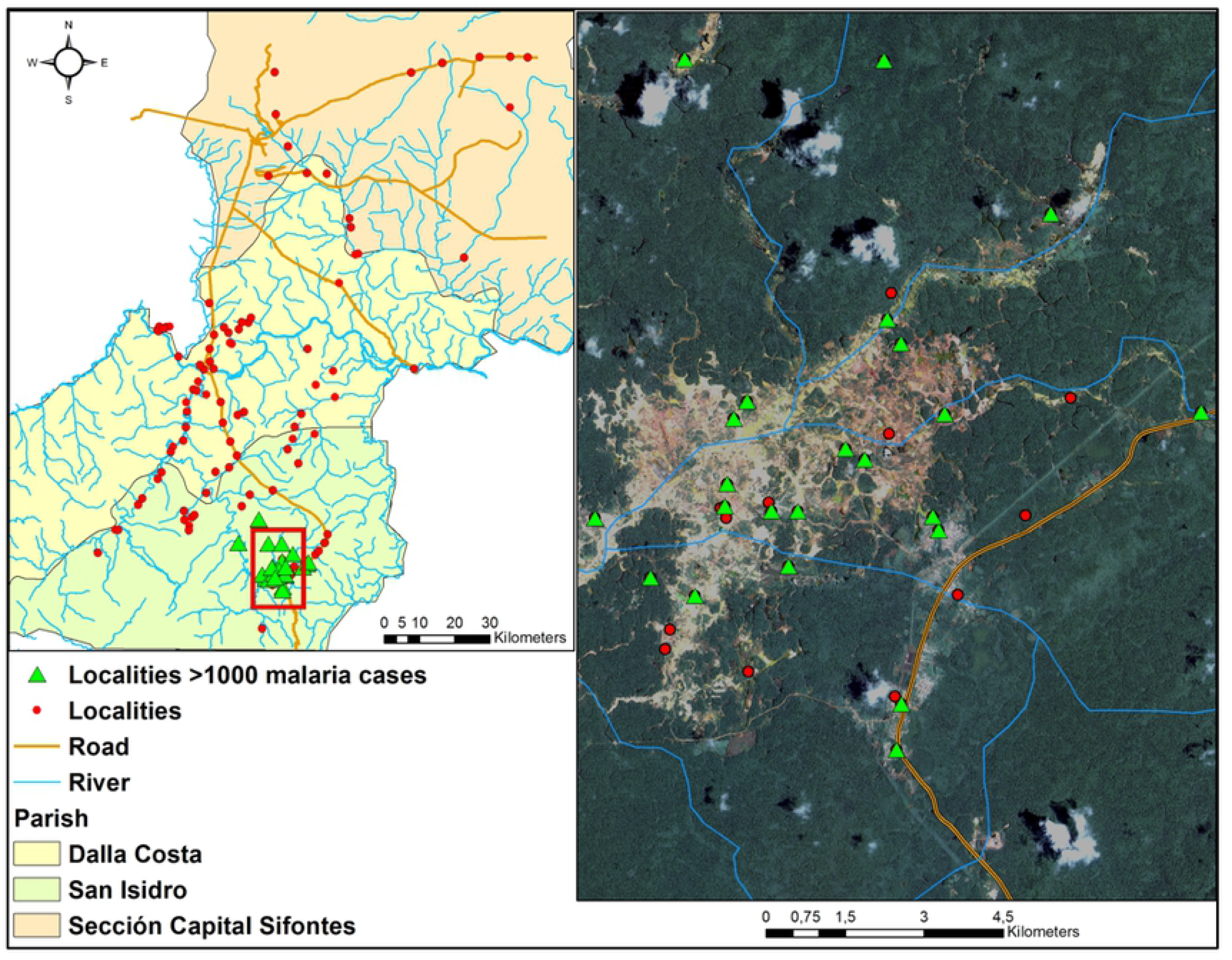
Left: Sifontes Municipality and its three parishes showing spatial distribution of all localities and some features of the general landscape. Right: satellite image (source: Landsat 8) of “Las Claritas” (framed in red window in San Isidro Parish); green triangles: localities with > 1,000 malaria cases across the study period in the deforested mining area; red dots are localities with < 1,000 malaria cases.

**Figure 8.**
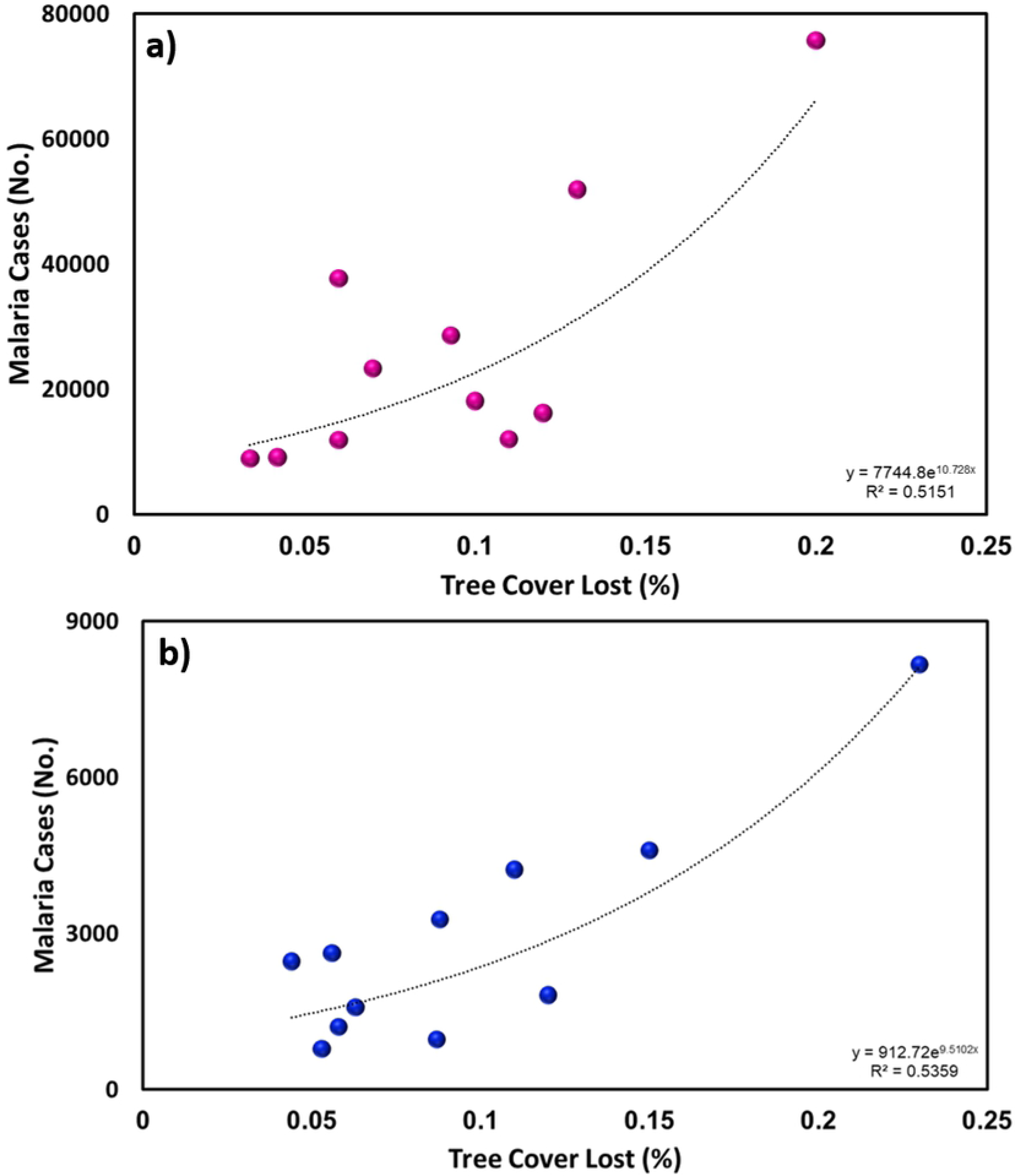
Percentage of deforestation (tree cover lost) and accumulated malaria (*P. vivax* + *P. falciparum* cases) in Sifontes Municipality (a) and Gran Sabana Municipality (b) in Bolívar State.

**Figure 9.**
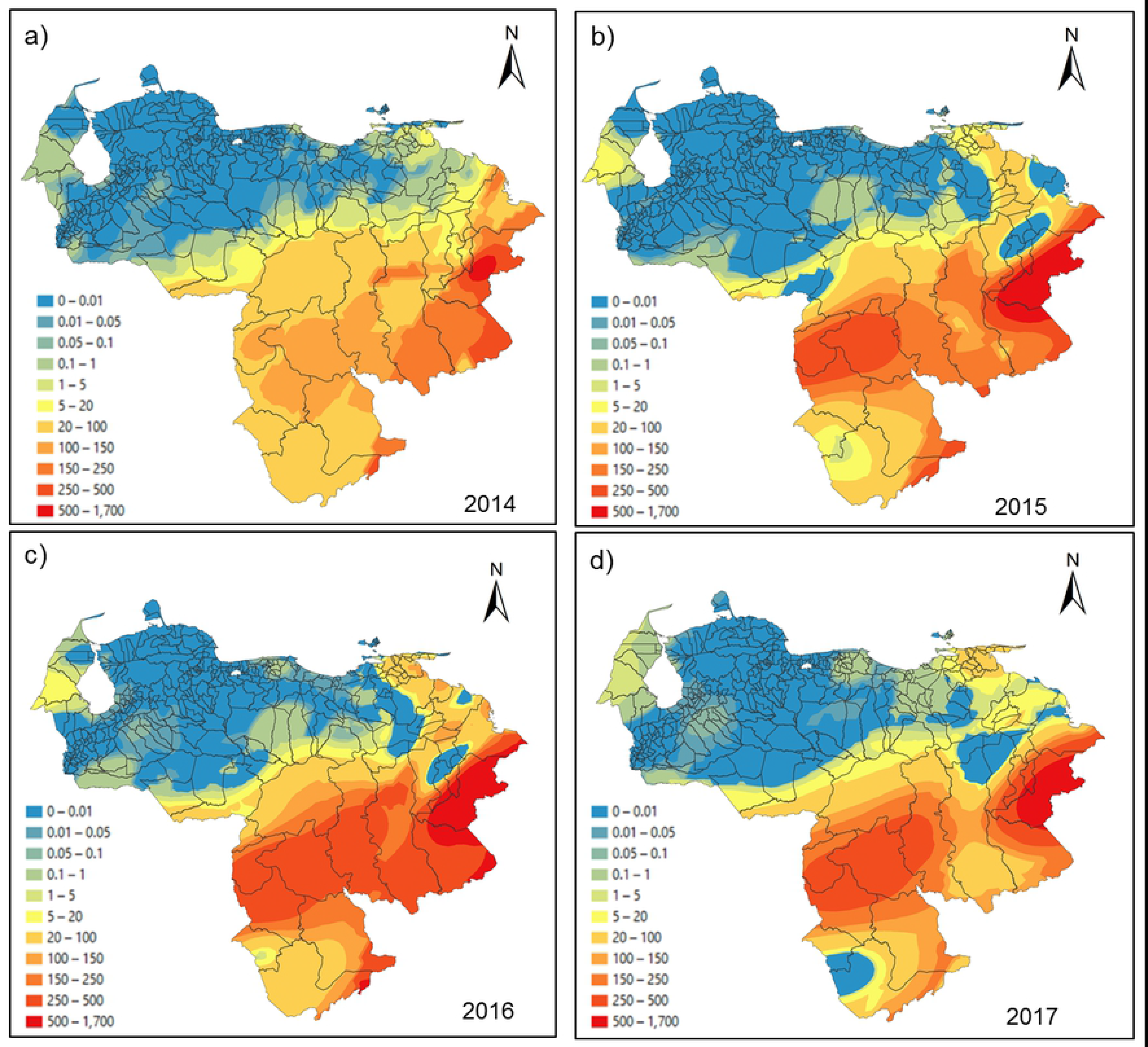
Spatial prediction maps (Venezuela malaria risk) for the 2014 (a), 2015 (b), 2016 (c) and 2017 (d) years derived from the ordinary gaussian kriging interpolation model. Note Year 2017, data available only through Epidemiology Week 21.

## Discussion

Malaria risk in southeastern Venezuela varied widely with most malaria cases reported in the mid-eastern and southern parishes of Bolívar state, where we identified two persistent hotspots. Local transmission in these infectious disease pockets accounted for most malaria transmission in the entire region (~ 61%) and country (>60%) over time (2007-2017). Both hotspots have been a long-standing regional source of *P. vivax* and *P. falciparum* transmission, as earlier studies suggested [7,13,19]. Our results support previous findings from Venezuela, Brazil and Peru showing that *P. vivax* malaria spatial heterogeneity is characterized by localities of high risk interspersed with others of moderate or low risk [11,30,31,32]. Due to their stability or persistence over time, the hotspots detected here could be predictive of prospective malaria incidence in the surrounding areas as has been found in other studies [33,34,35].

Our results show that disease patterns at larger spatial scales are driven by factors acting at local scales [36] such as mosquito ecology (especially larval habitats and host-seeking behavior) and at-risk human population dynamics (e.g., density, distribution, and mobility). In particular, our findings support the hypothesis that illegal gold mining is one of the leading local socioeconomic malaria drivers in southeastern Venezuela and a major factor explaining the malaria surge in the last years. First, we found *Plasmodium* cases clustered in areas deforested for gold mining activities. Secondly, both *P. vivax* and *P. falciparum* increased in incidence (4-8-fold) over time in those regions with a concomitant decrease of vegetation cover (3-6-fold) from mining activities. Finally, gold mining was significantly associated with a higher risk of acquiring malaria versus other occupations, with further analyses confirming occupational exposure of people to the risk of malaria infection regarding gender (male) and age groups (21-40 years). Illegal gold mining and the associated deforestation have rapidly increased and expanded in southern Venezuela since 2009 [37], especially in Sifontes municipality, Bolívar state.

Changes in land cover patterns resulting from deforestation can promote the emergence of larval habitats for these *Nyssorhynchus* (also known as *Anopheles*) vectors, thereby increasing mosquito abundance [38], vector-host contact [39] and consequently transmission risk [40], especially in human settlements located near the forest fringe. Also, the forest patches generated by fragmentation can provide resting sites and shaded refuges for adult mosquitoes, increasing their survival [41]. In the study areas, *Ny*. *albitarsis* s.l. (larvae develop in open lagoons and herbaceous swamps) and *Ny*. *darlingi* (found mostly on river margins) co-occur in forest lagoons [42,43]. Thus, we hypothesize that forest fragmentation by mining activities constitutes a habitat gain for both species. Earlier studies indicated that the most productive breeding site types for *Ny. albitarsis* s.l. and to a lesser degree for *Ny. darlingi*, are abandoned open lagoons or mining dug-outs left after clearing vegetation [42,43]. Preliminary research has also found that high malaria case localities in the epicenter of illegal mining operations have a higher proportion of breeding sites positive for both vector species within 1 km of their epicenters compared with low malaria localities (unpublished data). Herein we suggest that the identification and quantification of ecological mechanisms that increase host-vector contacts in mining settlements in southeastern Venezuela should be a priority for regional malaria control.

The estimated values of *PvR_0_* and *PfR_0_* in San Isidro and Dalla Costa were ~2, meaning that one single human infected with *P. vivax* or *P*. *falciparum* from a mosquito bite was able to produce up to 2 secondary malaria cases during his or her entire infectious period. These unexpectedly low figures are in contrast to the high annual parasite incidence for both parasite species reported (e.g., 4,672 cases per 1,000 inhabitants) and the high *Plasmodium* Infectivity Rate (IR) detected for *Ny. albitarsis* s.l. (5.4%: 27/499) and *Ny. darlingi* (4.0%: 37/926) from collections (2009–2012) in this area [20]. Indeed, the work of these latter authors emphasizes that in this gold mining area, *Ny. albitarsis* s.l. increased its *Plasmodium* infection rate more than 3 times in ten years [17]. Although our approach is useful in providing a preliminary estimate of *R_0_*, it is likely an underestimate for four reasons: first, stochastic fluctuations in the early stages of the endemic-epidemic curve obscure the measure of the intrinsic growth rate, *r* [26]; second, reporting inaccuracies due to poor case management due to limited access to health services, asymptomatic infections particularly in *P. vivax* [7,44] very likely bias the current incidence data; third, incubation period duration in both parasites, the number of relapses and relapse patterns for *P. vivax* (to calculate the serial interval, *V)* in southern Venezuela remain to be determined. Finally, even when the relationship between *r* and *R_0_* can be measured with some confidence, it is highly dependent on other entomological and parasitological parameters that we were not able to account for here, such as vectorial capacity and the recovery rate of humans from parasitemia. Malaria transmission should ideally be measured in a way that accounts for all components of the parasite lifecycle [45]. A recent study carried out in the Brazilian Amazon region reported *R_0_* of *P. vivax* transmitted by *An. darlingi* ranging from 3.3 to 58.7, suggesting malaria hyperendemicity at levels similar to those found for *P. falciparum* in sub-Saharan Africa [46]. Had we included in the *R_o_* estimate the vectorial capacity of the two main biting species in southeastern Venezuela, *Ny. darlingi* and *Ny. albitarsis* s.l, we would have expected values similar to those determined for Amazonian Brazil.

The malaria epidemic in the country has been fueled by financial constraints for procurement of malaria control commodities (such as insecticides, drugs, diagnostic supplies, and mosquito nets) and surveillance activities, and lack of provision and implementation of services [6,47]. In particular, the Venezuelan economic crisis has promoted the informal sector, with illegal gold mining featuring as one of the country’s fastest-growing shadow economic activities [48]. As a result, migration within the country has increased toward southeastern Venezuela (Sifontes Municipality) where gold mining activities are clustered and where malaria hotspots have been identified. This highly mobile human population of miners, predominantly young males migrating in search of economic gain, work and sleep in or near the forest (forest fringe). They often live in provisional camps with incomplete walls that promote even higher intensities of mosquito-host contact [4,5]. Many such economic migrants reintroduce *Plasmodium* to previously controlled areas where competent vectors are abundant. Concomitantly, the dismantling of the epidemiological surveillance systems at the national level has unintentionally prompted the reappearance and spread of malaria in regions that have not experienced autochthonous transmission in decades [7].

The spatial connectedness of malaria foci, related to travel behavior of (infected) individuals, has been used to identify so-called sources and sinks of *Plasmodium* within a transmission network approach [49]. Sources (originating and exporting cases) and sinks (receiving imported cases) of malaria parasites mostly occur in epidemiological settings when humans travel frequently between locations characterized by substantial heterogeneity in malaria transmission [15,50]. Consequently, we hypothesize that a source-sink malaria parasite (metapopulation) dynamic accounts for the current spatial distribution and persistence of this infection in Venezuela. As we showed, since 2014, malaria has spread from the south and northeastern coastal regions to the lowland central savannas and west Andes piedmont ecoregions where malaria had previously been eliminated, increasing the population at-risk to around 50% compared to 34.4% in 2010. Our results also confirm the importance of malaria linked to mobile illegal gold miners and their role in its spread among the Guiana Shield countries [51,52,53,54]. Understanding the spatiotemporal variability between *Plasmodium* and human movement (spatial demography) will be key in designing and implementing a strategic malaria control program in Venezuela.

The rapidly increasing malaria burden in Venezuela is affecting neighboring countries, particularly Brazil and Colombia [7]. Migration to Brazil occurs via Route 10, the sole highway that links southern Venezuela and northern Brazil. Unfortunately, this road runs through the hotspots of San Isidro and Dalla Costa. Infected people crossing these borders complicate each country’s malaria program by the potential reintroduction of the disease in areas where it had previously been reduced or eliminated; enhancing malaria transmission near border areas; promoting case spillover, and possibly spreading drug resistance alleles across Brazil. Another important risk includes the effects of inadequate treatment that can increase malaria-related mortality. According to the Brazilian Ministry of Health, malaria cases imported from Venezuela into neighbouring Roraima State, Brazil have increased from 1,538 (2014) to 4,478 (2018), representing up to 85% of the reported malaria cases of that border state [7,23].

Mutations linked to drug-resistant in *P. falciparum* in Venezuela remains poorly understood, although surveillance for resistance markers is now recommended in some Guiana Shield countries such as Guyana and in the mining areas where artemisinin is available for sale and self-treatment is common [55]. Considering that there are already multi-drug resistant *Plasmodium falciparum* lineages against common drugs such as Chloroquine and SP circulating in Venezuela [55,56], and the recent report of novel mutations linked to the delay of artemisinin efficacy in Guyana [16], it is possible that multi-drug resistant parasite strains that also include artemisinin delayed mutations, could start to circulate in the southern Venezuela hotspots.

## Conclusions

We found evidence that gold-mining is driving malaria hotspots in Venezuela and those high transmission pockets were critical in the surge of malaria observed since 2014 onwards. These gold-mining areas likely make malaria transmission resilient to interventions, as they not only sustain transmission but also can restore it (“rescue effect”) after interventions have reduced malaria locally or even achieved local elimination in other areas as has been observed in Venezuela [15,57]. Transmission in such settings is considered the Gordian knot for achieving malaria elimination in Latin America [5,58,59]. Hotspot-targeted control has been hypothesized as an effective approach to reduce the burden of malaria in areas of heterogeneous malaria transmission [57,60]. Thus, a program focused on rapid diagnosis and timely treatment, vector control, and monitoring for drug/insecticide resistance is urgent and essential in these hotspots. Increased enforcement of the malaria control program in the rest of Venezuela is pivotal to lower the risks of reintroduction to vulnerable areas. Given the current context, successful control of the ongoing malaria epidemic in Venezuela requires not only national but regional coordination, as evidenced by the cross-border malaria spillover. Without international coordinated efforts, the progress achieved toward malaria elimination in Latin America over the past 18 years could be easily reversed.

## Supporting Information

S1 Table (Supplementary Material)

S1 Figure (Supplementary Material)

## Acknowledgements

We acknowledge the logistic support (field and laboratory) provided by Angela Martinez, Porfirio Acevedo, Mayida El Souki, Virginia Behm and Nelson Moncada. This work was supported by the Council for Sciences and Humanities Development (Grant CDCH-PG-0382182011 to MEG and JEM) and partially funded by the National Institutes of Health, USA, Grant R01 AI110112 to JEC.

## Author Contributions

**Conceptualization:** Maria E Grillet

**Data curation:** Maria E Grillet, Juan V Hernández-Villena, Maria F. Vincenti-Gonzalez

**Formal analysis:** Maria E Grillet, Juan V Hernández-Villena, Maria F. Vincenti-Gonzalez

**Funding acquisition:** Maria E Grillet, Jorge E Moreno, Jan E Conn

**Investigation:** Maria E Grillet, Jorge E Moreno, Jan E Conn, Juan V Hernández-Villena

**Methodology:** Maria E Grillet, Juan V Hernández-Villena, Maria F. Vincenti-Gonzalez

**Project administration:** Maria E Grillet

**Resources:** Maria E Grillet, Jan E Conn, Ananias A Escalante, Jorge E Moreno, Alberto-Paniz Mondolfi, Oscar Noya, Rachel Lowe, Martin Llewellyn, Adriana Tami

**Software:** Maria E Grillet, Juan V Hernández-Villena, Maria F. Vincenti-Gonzalez

**Supervision:** Maria E Grillet

**Validation:** Maria E Grillet, Jorge E Moreno

**Visualization:** Maria E Grillet, Juan V Hernández-Villena, Maria F. Vincenti-Gonzalez

**Writing – original draft:** Maria E Grillet, Jan E Conn, Ananias A Escalante

**Writing – review & editing:** Maria E Grillet, Jan E Conn, Ananias A Escalante, Jorge E Moreno, Alberto-Paniz Mondolfi, Oscar Noya, Rachel Lowe, Martin Llewellyn, Adriana Tami

---

**S1 Figure.** Age distribution of *P. vivax* (upper panel: a, b) and *P. falciparum* (lower panel: c, d) among malaria patients of San Isidro (left side: a, c) and Dalla Costa (right side: b, d) parishes (northeastern of Bolivar state) across the study period.

**S1 Table**. Ro estimations for *Plasmodium vivax* and *Plasmodium falciparum* for San Isidro and Dalla Costa malaria cases in the Bolivar state (South-eastern Venezuela) during the 2014-2017 period.

